# Gut microbiota dysbiosis induced by brain tumor modulates the efficacy of immunotherapy

**DOI:** 10.1101/2024.08.18.608488

**Authors:** Hyeon Cheol Kim, Hyun-Jin Kim, Jeongwoo La, Won Hyung Park, Sang Hee Park, Byeong Hoon Kang, Yumin Kim, Heung Kyu Lee

## Abstract

**Background:** Although the influence of gut microbiota on various tumors has gained recognition, the intricate mechanisms underlying their effects on brain tumors remain largely unexplored. In this study, our objective was to unveil pivotal gut microbiota that contribute significantly to the anti-tumor immune response against brain tumors.

**Results:** During the progression of brain tumors, our research uncovered a notable shift in the gut microbiome, as revealed through comprehensive 16S rRNA sequencing analysis. This shift coincided with a substantial decrease in the concentration of tryptophan in fecal samples, indicating a potential link between tryptophan levels and microbiome composition. Intriguingly, dietary supplementation with tryptophan significantly mitigated these microbial alterations, suggesting a restorative effect on the gut microbiota composition. This intervention not only reversed the changes observed in the gut microbiome but also markedly improved survival rates, a phenomenon that was determined to be dependent on the gut microbiota. The mechanism underlying this improvement appears to involve an enhancement in T cell circulation, which in turn boosts the efficacy of immunotherapeutic approaches. Upon further investigation into the specific microbial species that were positively influenced by tryptophan supplementation, some candidates were screened. Among several gut microbiota strains restored by tryptophan supplementation, the most significant is *Duncaniella dubosii*. Through sole colonization into germ-free mice, we found that the presence of *Duncaniella dubosi*i alone was able to replicate the beneficial effects observed with tryptophan supplementation.

**Conclusions:** This study highlights *Duncaniella dubosii*’s crucial role in linking tryptophan supplementation to positive shifts in the gut microbiome, immune modulation, and enhanced survival during brain tumor progression. It underscores the complex interaction between diet, microbiota, and immune responses, offering novel insights for boosting cancer immunotherapy’s success.

## Background

Glioblastoma (GBM), known for its dire prognosis [1], remains one of the most challenging brain tumors to treat, despite extensive research efforts and the availability of various anticancer therapies. Unlike some other tumor types, survival rates for GBM have seen limited improvement [2]. While the brain is no longer considered an immune-privileged tissue [3, 4], GBM’s poor response to treatments stems from its unique characteristics, notably the scarcity of highly cytotoxic effector cells within the tumor microenvironment [5]. This deficiency hampers the effectiveness of immunotherapies targeting these cells [6, 7]. Moreover, the observation of T cell sequestration in the bone marrow in GBM cases exacerbates treatment challenges [8].

Nonetheless, leveraging T cells against GBM remains a promising strategy. Given the tumor’s heterogeneity [9, 10], third-generation therapeutic approaches are likely to offer improved outcomes compared to earlier iterations. Additionally, the diverse repertoire of T cell receptors allows for targeting a wide array of tumor antigens, and the widespread circulation of T cells throughout the body presents opportunities for effective treatment. Further research is imperative to fully comprehend and harness the potential of T cells in the context of GBM.

Commensal microbiota, primarily found in the skin and gut, play a vital role in maintaining host health and impacting cancer development, notably in gastrointestinal tumors, where they operate through diverse mechanisms [11–13] and affect treatment outcomes [14–17]. However, their influence on cancers originating in distant tissues [18–20], like brain tumors, is less explored. Unlike in neurodegenerative and mood disorders, where the gut-brain axis receives significant attention, the precise mechanisms through which gut microbiota influence health outcomes are not fully understood, particularly in GBM [21]. Despite growing interest, comprehensive documentation regarding the role of gut microbiota in mediating anti-tumor immune responses in GBM remains limited.

Here, we determined the impact of the gut microbiota on the antitumor response in the brain. Using GBM animal models, we found that a stable gut microbiota is critical for the host defense response in brain tumor progression. We used tryptophan (Trp) supplementation to restore a disrupted gut flora that resulted from tumor progression and found an enhanced antitumor cytotoxic T lymphocyte (CTL) response and increased efficacy of immunotherapy via improved T cell circulation. Importantly, these effects depended on the gut microbiota and a particular bacterial strain that closely mimicked the positive effects of Trp supplementation.

## Results

### Homeostatic gut microbiota plays a crucial role in defending against brain tumor

To investigate the importance of gut microbiota in brain tumors, we analyzed changes in gut microbiota across two mouse models (Additional file 1: Fig. S1A–D). Both orthotopic implantation and genetically engineered GBM models exhibited similar alterations. Initially, we evaluated microbial diversity [22], finding no significant changes in several alpha diversity indices (Additional file 1: Fig. S1E–H). However, principal coordinate analysis (PCoA) revealed a distinct shift in beta diversity, indicating gut microbiota clustering according to tumor progression (Fig. 1A, B). This suggests concurrent changes in gut microbiota during brain tumor advancement. Additionally, microbial composition showed similar shifts at the phylum level, with fewer Bacteroidetes and more Firmicutes (Fig. 1C, D), alongside notable alterations in specific species across both models (Fig. 1E, F). These observations signify a shift in gut microbiota accompanying brain tumor progression.

**Fig. 1.**
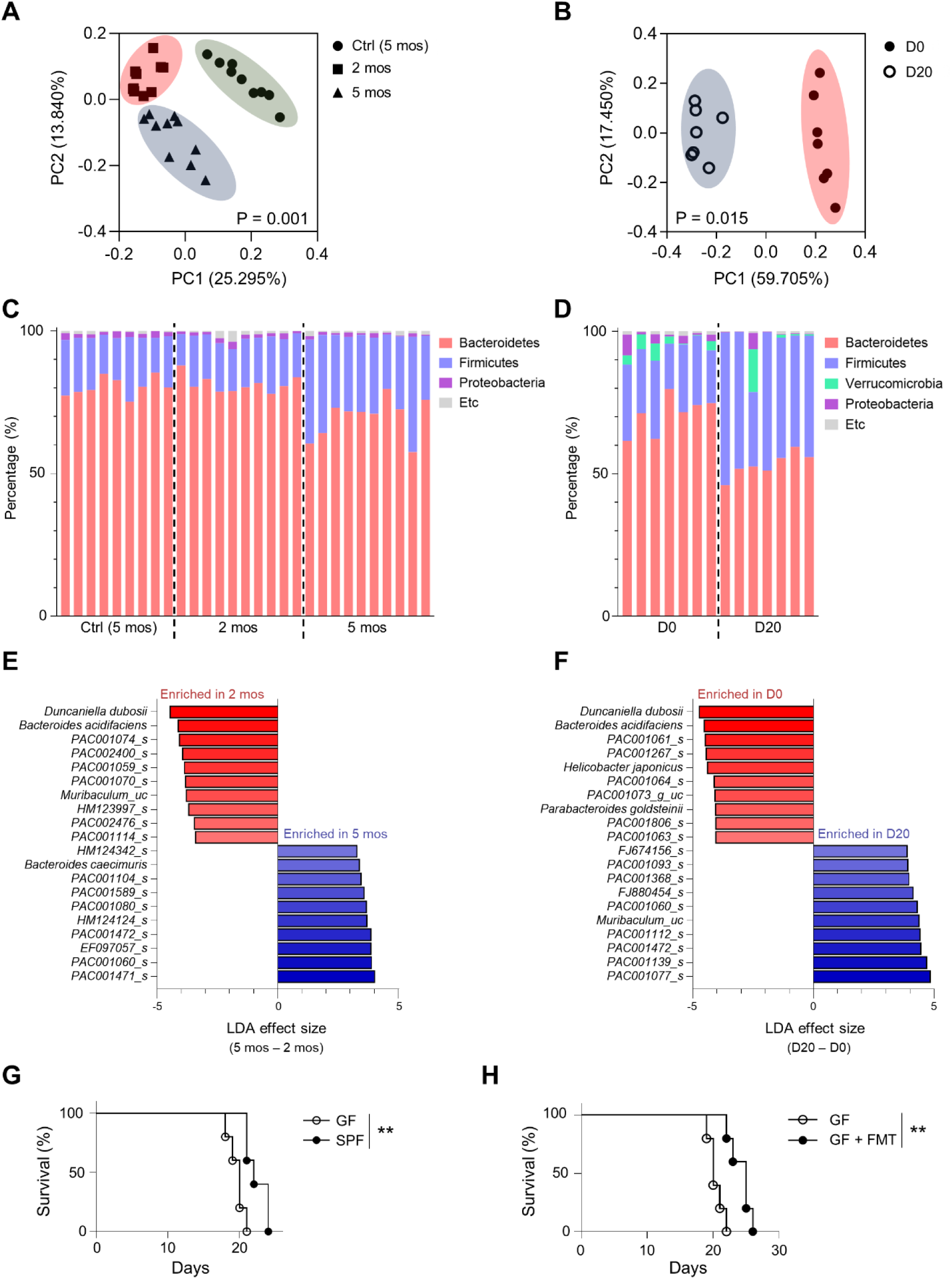
The homeostatic gut microbiota is important for the defense against brain tumors. **A**-**F** We performed 16S rRNA sequencing analysis on feces collected from genetically engineered GBM mice at 2 months (2 mos) and 5 months (5 mos) after tumor induction (**A**, **C**, **E**) and from orthotopic implantation mice at 20 days (D20) after tumor implantation (**B**, **D**, **F**). Control mice were following as: genetically engineered model, 5 mos; orthotopic implantation model, D0. (**A**) Ctrl (5 mos) (n = 9), 2 mos (n = 10) and 5 mos (n = 10) (**B**) D0 (n = 7) and D20 (n = 7). (**A**, **B**) During brain tumor progression, beta diversity of gut microbiota was compared using PCoA analysis. (**C**, **D**) Phylum level comparison of gut microbial composition. Phyla present at less than 1% are designated as etc. (**E**, **F**) Linear discriminant analysis effect size (LEfSe) was used to identify the top 10 OTUs that increased or decreased at the species level, and these were selected as the notably changed gut microbiota (*P* < 0.05). **G** Survival of mice after transplantation of GL261 tumor cells into germ-free (GF) mice (n = 5) and specific pathogen-free (SPF) mice (n = 5). **H** Survival of tumor-bearing GF mice after fecal microbiota transplantation (FMT) from healthy SPF mice (GF + FMT, n = 5) vs. control GF mice (n = 5). Beta diversity (**A**, **B**) was calculated based on generalized Uni-Frac algorithms. Survival data (**G**, **H**) was analyzed using a log-rank test. \**\*P <* 0.01

Based on previous findings, we determined that gut microbiota changes accompanying brain tumor progression are consistent across both models. We hypothesized these changes may represent survival strategies of tumors, leading to the hypothesis that they could impact the health of host bearing brain tumors. Survival comparisons between GF and SPF mice with brain tumors showed longer survival in SPF mice (Fig. 1G). However, the presence or absence of commensal microbes affects various factors, including immune system development [23]. To assess commensal microbe impact under equivalent immune conditions, we transplanted SPF mouse fecal microbiota to GF mice, significantly improving survival in recipient group (Fig. 1H). Thus, homeostatic microbiota plays a critical role in host defense against brain tumors.

### Tryptophan supplementation restores the gut microbiota and enhances survival during brain tumor progression

Changes in gut microbiota may be influenced by the nutrient, particularly amino acids, in the gut [24, 25]. Thus, we hypothesized that alterations in gut microbiota could be linked to changes in amino acid levels. To investigate, we used mass spectrometry to quantify amino acids and observed a significant reduction in tryptophan levels in mouse feces (Fig. 2A) and serum (Additional file 1: Fig. S2A–D) during tumor progression. Furthermore, there was an increase in microbial metabolic processes consuming tryptophan (Additional file 1: Fig. S2E–H) and a decrease in processes producing tryptophan (Additional file 1: Fig. S2I–L). These findings suggest a quantitative reduction in tryptophan levels as GBM progresses. Considering the close relationship between gut microbiota changes and gut nutritional status, we explored the correlation between tryptophan reduction in brain tumor conditions and alterations in gut microbiota.

**Fig. 2.**
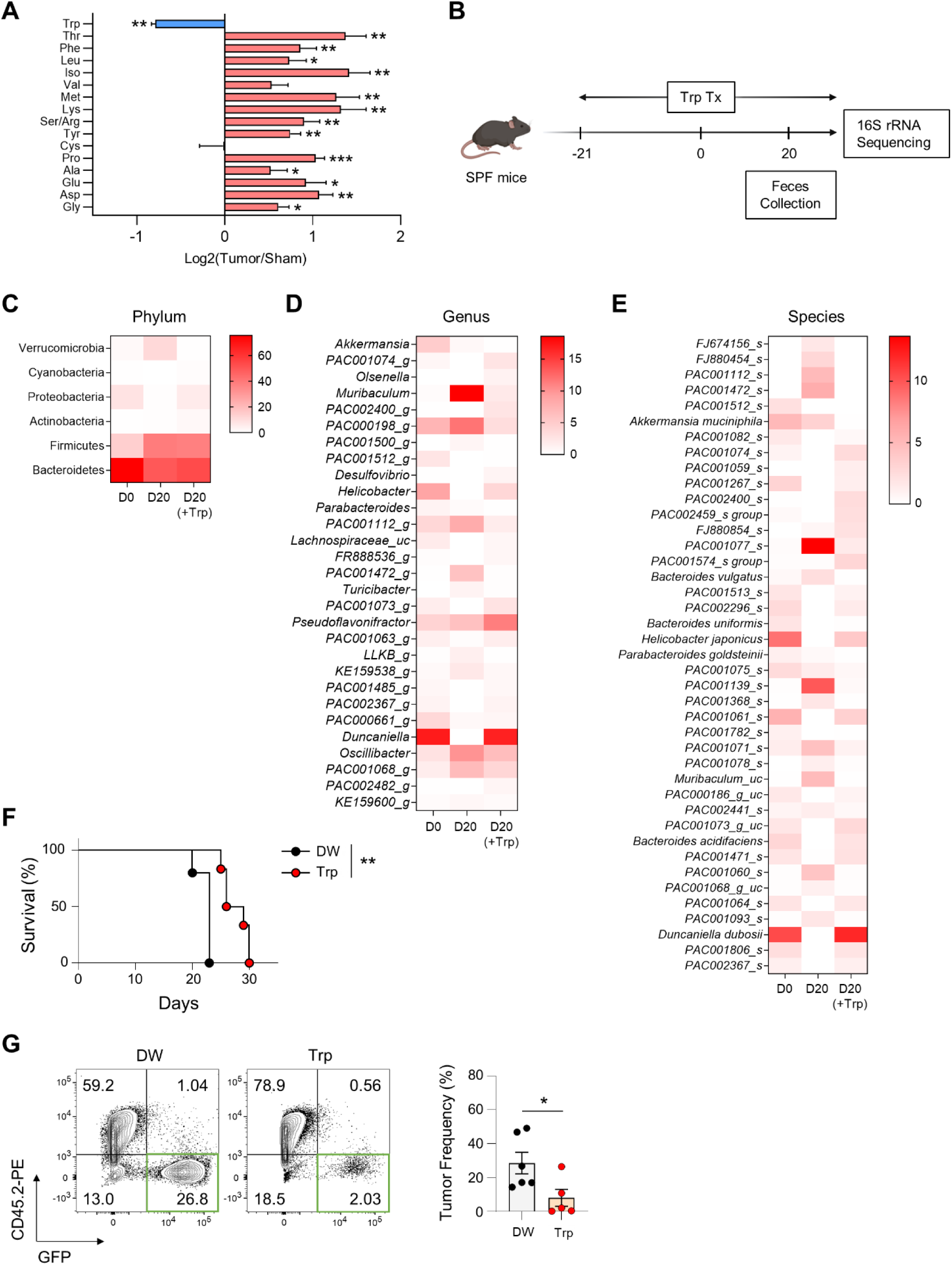
Tryptophan supplementation reversed tumor-induced gut microbiota changes and improved brain tumor survival. **A** Amino acids were measured by mass spectrometry in feces collected on D20 after tumor (n = 5) or sham (n = 5) injection into SPF mice. **B**-**E** Gut microbial abundance heatmap at D0 vs. D20 vs. D20 (+Trp). **B** Three weeks before tumor implantation, mice received supplemental Trp in the drinking water. 20 days after tumor injection, feces were collected from mice supplemented with Trp (n = 6) or with distilled water (DW, n = 5), and the fecal microbiota were compared with the control sample (D0, n = 5) using 16S rRNA sequencing. The gut microbial abundance is illustrated on the heatmap at the phylum (**C**), genus (**D**), and species (**E**) levels. **F** Survival of mice with a brain tumor was compared in the control group (DW, n = 5) vs. the Trp group. Trp treatment was the same as in (**B**). **G** For tumor growth comparison, GL261-GFP cells were transplanted into mice supplemented with Trp or with DW. Statistical significance was determined by a two-tailed Student’s *t*-test in (**A**, **G**), a two-way ANOVA test in (**C**-**E**), and a log-rank test in (**F**). Survival was compared in four or more independent experiments. Data are presented as mean ± standard error mean (s.e.m.) *\*P <* 0.05, \**\*P <* 0.01

We aimed to restore the gut microbiota altered during tumor progression. Amino acids significantly influence gut microbiota composition [25]. Given the decreased tryptophan (Trp) levels during tumor progression, we implemented dietary supplementation of Trp in mice, which rescued microbial changes induced by tumor progression (Fig. 2B). While no significant difference was observed at the phylum level (Fig. 2C), Trp supplementation showed a trend towards a composition resembling a homeostatic gut microbiota at the genus and species levels. Particularly, the *Duncaniella* genus and *Duncaniella dubosii* strain exhibited the most significant differences in Trp-treated mice (Fig. 1E, F and Fig. 2D, E). Tumor-bearing mice treated with tryptophan showed significant improvements in survival (Fig. 2F) and reductions in tumor burden (Fig. 2G). Thus, Trp supplementation restored gut microbiota homeostasis and improved survival by reducing tumor burden.

### Trp treatment leads to an increase in CD8 T cell circulation, thus enhancing the antitumor response and synergizing with anti-PD-1 therapy

Based on our findings, Trp treatment was expected to enhance the antitumor response. Transcriptomic analysis showed increased CD8 T cell cytotoxicity in the Trp-treated group (Additional file 1: Fig. S3A, B and Fig. 3A, B). This was confirmed by elevated cytotoxicity in CD8 T cells from Trp-supplemented tumor-bearing mice (Fig. 3C). Additionally, the Trp group exhibited higher expression of effector molecules, with comparable exhaustion molecule expression (Additional file 1: Fig. S3C–H and Fig. S4). CD8 T cell depletion experiments revealed reduced survival in Trp-treated conditions (Fig. 3D), contrasting with conventional CD8 T cell depletion alone [26]. Survival did not improve in athymic mice despite Trp supplementation (Fig. 3E). Thus, the enhanced CTL response induced by Trp likely contributes to improved survival.

**Fig. 3.**
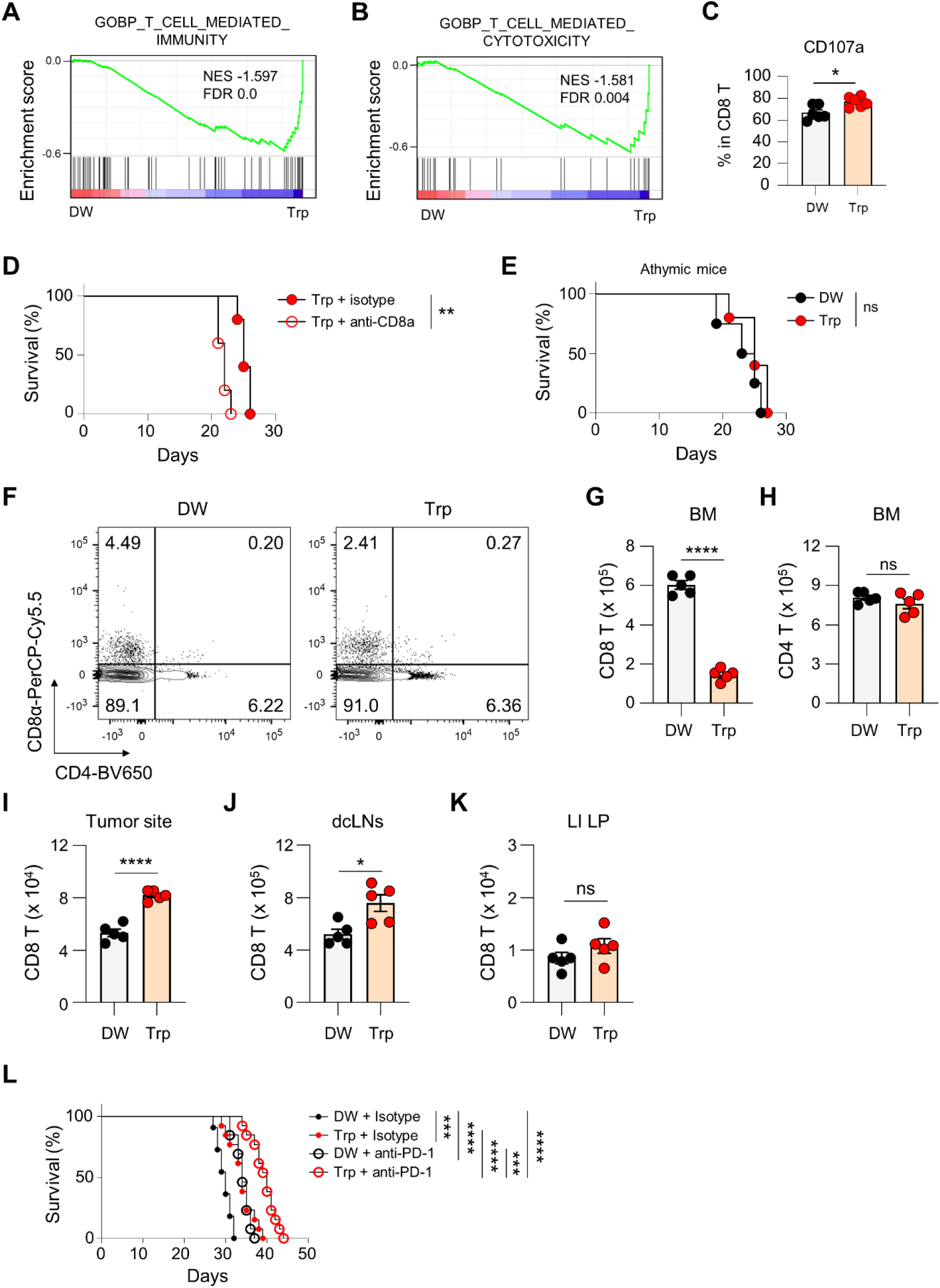
Tryptophan treatment increased CD8 T-cell circulation, thus increasing the antitumor response and enhancing anti-PD-1 therapy. **A**, **B** Gene Set Enrichment Analysis (GSEA) using gene set (**A**) GOBP_T_CELL_MEDIATED_IMMUNITY and (**B**) GOBP_T_CELL_MEDIATED_CYTOTOXICITY in the tumor-infiltrating CD8 T-cell subset. The CD8 T cells of the control group (DW) were compared to the CD8 T cells of the Trp experimental group (Trp). The transcriptomes of immune cells that were sorted from the brains of tumor-bearing mice on D20 after tumor transplantation were determined by scRNA-seq. **C** 20 days after tumor transplantation, CD107a expression was measured by flow cytometry in tumor-infiltrating CTLs from control and Trp-treated tumor-bearing mice. **D** In tumor-bearing mice, survival was compared for mice in which CD8a+ cells were depleted by intraperitoneal injections of an anti-CD8a antibody vs. mice injected with an isotype antibody, using two or more independent experiments. **E** In tumor-bearing mice, survival was compared for athymic mice with (Trp) or without Trp supplementation (DW), as described in Fig. 2B. **F**-**K** On D20 after tumor transplantation for DW and Trp groups, CD8 and CD4 T cells were quantified by flow cytometry in bone marrow (**F**-**H**). Similarly, CD8 T cells were measured in the brain (**I**), deep cervical lymph nodes (**J**), and colon lamina propria (**K**). **L** For immunotherapy, anti-PD-1 antibody was injected intraperitoneally. Survival was compared for the DW + isotype (n = 11), Trp + isotype (n = 13), DW + anti-PD-1 (n = 13), and Trp + anti-PD-1 (n = 13). Statistical significance was calculated using a two-tailed unpaired Student’s *t*-test in (**C**, **G**-**K**) or a log-rank test in (**D**, **E**, **L**). Data are presented as mean ± s.e.m. *\*P <* 0.05, \**\*P <* 0.01, \*\**\*P <* 0.001, \*\*\**\*P <*0.0001, ns: not significant.

Nevertheless, CD8 T cell numbers remained similar at d20 (Additional file 1: Fig. S3I–K). Integrative analysis comparing d20 samples with normal or earlier brain tumor microenvironment (TME) samples revealed significant T cell infiltration by day 10 (Additional file 1: Fig. S5, GSE201559 and GSE200533). T cells were confined to the bone marrow between days 10 and 20 post-tumor transplantation (Additional file 1: Fig. S6A–C). Bone marrow analysis on day 15 showed fewer CD8 T cells in Trp-treated group (Fig. 3F–H), while brain, draining lymph nodes, and gut had more CD8 T cells in Trp-supplemented mice (Fig. 3I-K). Despite unchanged brain tissue T cell infiltration due to blood-brain barrier compromise from tumor growth, systemic circulation varied (Additional file 1: Fig. S6D–G). Trp supplementation possibly enhanced T cell circulation, overcoming bone marrow sequestration and boosting immunotherapy efficacy. Indeed, Trp supplementation boosted anti-PD-1 treated mice survival (Fig. 3L).

### Among gut microbiota, particularly *D. dubosii* is crucial in Trp-driven enhancement of survival

Trp deficiency in TME leads to immune dysfunction and increased apoptosis [24]. Brain Trp levels in tumor-bearing mice were comparable to controls (Fig. 4A). *Eif2ak4* (Gcn2) expression, responsive to amino acid deficiency [27], remained unchanged with Trp supplementation (Additional file 1: Fig. S7A, B). Apoptosis in T and B cells within TME showed no significant difference regardless of dietary Trp (Additional file 1: Fig. S7C, D). However, fecal Trp levels were higher in the Trp-treated group (Fig. 4B). GF mice treated with Trp did not show improved survival (Fig. 4C), but FMT from Trp-treated SPF mice enhanced survival (Additional file 1: Fig. S7E), highlighting gut microbiota’s crucial role in Trp-mediated survival enhancement. Since gut barrier integrity remained unchanged during tumor progression or Trp treatment (Additional file 1: Fig. S7F), we conclude that Trp-mediated effects rely on gut microbiota without involving translocation.

**Fig. 4.**
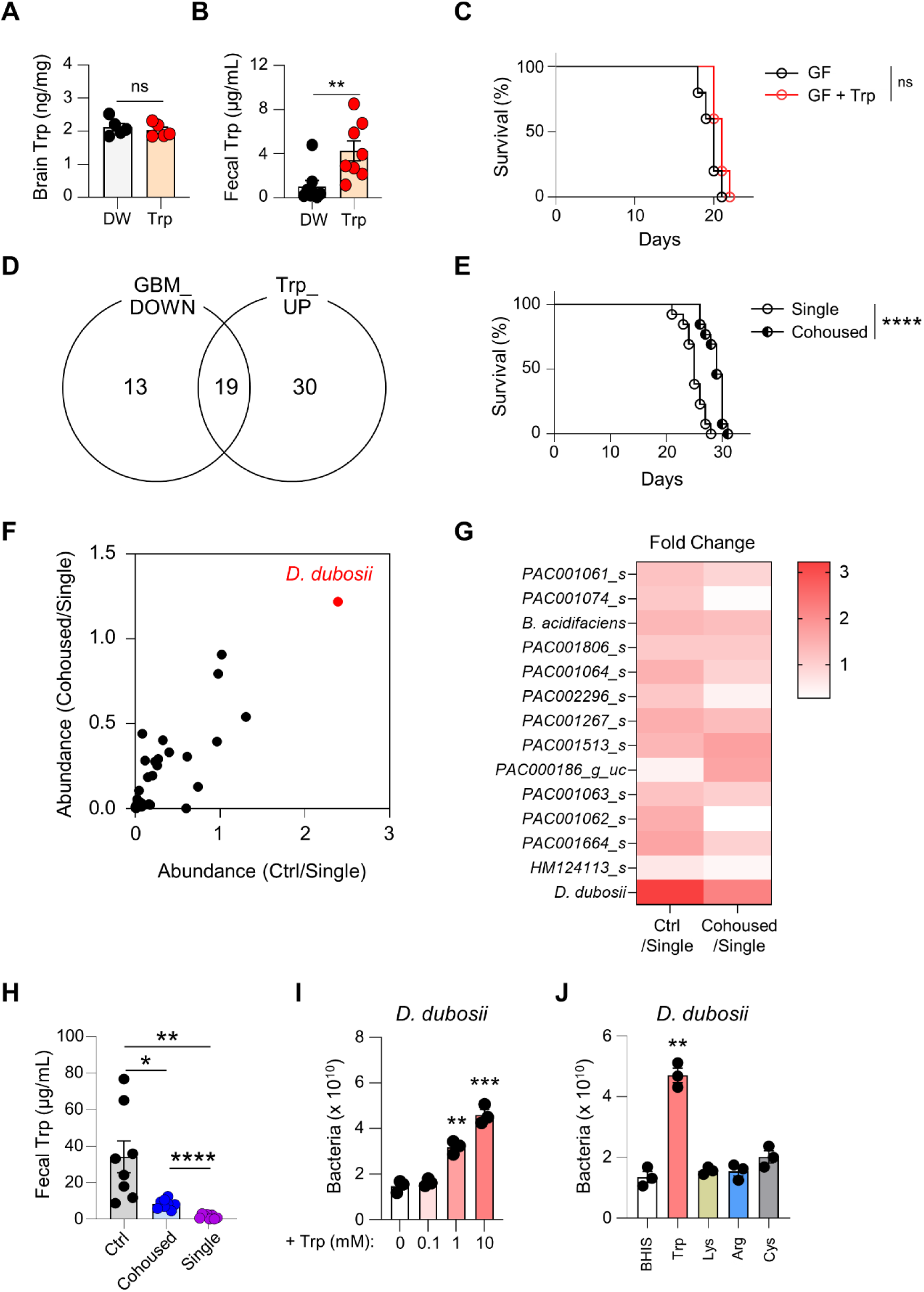
Trp enhances antitumor response in brain tumors through gut microbiota changes, especially *D. dubosii*. **A**, **B** 20 days after tumor transplantation, the amount of Trp was measured in brain tissue (**A**) (n *=* 5 each group) and feces (**B**) (n *=* 8 for each group) through ELISA. **C** Survival of tumor-bearing GF mice supplemented with Trp was determined (n *=* 5 for each group). **D** The gut microbiota strains that decreased during brain tumor progression and increased after Trp supplementation in tumor-bearing mice were identified in a Venn diagram. All microbial species changes were significant (*P* < 0.05). **E** Survival of tumor-bearing mice co-housed with healthy mice (co-housed mice) compared with mice that were housed singly (single). Healthy mice (Ctrl). n = 13 for each group. **F**, **G** Microbial abundance ratios were compared in an XY plot (**F**) and heatmap (**G**) for co-housed, single-housed, and healthy control mice. Dots represent the strains in (**D**), and the red dot is *D. dubosii*. **H** In the co-housing experiment, the amount of fecal Trp was measured using ELISA on day 20 (n *=* 8 for each group). **I**, **J** *D. dubosii* cells were counted after 3 days of growth in BHIS medium with various concentrations of Trp (**I**) or with other amino acids at 10 mM (**J**). Statistical significance was calculated using a two-tailed unpaired Student’s *t*-test in (**A**, **B**, **H**-**J**) or with a log-rank test in (**C**, **E**). Data are presented as mean ± s.e.m. *\*P <* 0.05, \**\*P <* 0.01, \*\**\*P <* 0.001, \*\*\**\*P <*0.0001, ns: not significant.

To pinpoint bacterial contributors to enhanced survival, we identified 19 microbiota strains declining during brain tumor progression yet increasing with Trp supplementation in tumor-bearing mice (Fig. 4D). Co-housing tumor-bearing mice with normal counterparts, which normalized gut microbiota, notably improved survival compared to individually housed mice (Fig. 4E). Among these strains, *D. dubosii*, declining during tumor progression and effectively transferred via co-housing, emerged as the most significant (Fig. 4F, G). Its abundance showed the most substantial change following Trp treatment, highlighting its pivotal role. Additionally, co-housed mice exhibited higher fecal Trp levels than singly-housed counterparts (Fig. 4H).

Next, we considered a potential causal relationship between the abundance of *D. dubosii* and the changes in Trp levels during brain tumor progression. We hypothesized that (1) Trp is essential for *D. dubosii* or (2) *D. dubosii* secretes Trp. Growth experiments using various concentrations of Trp revealed that the abundance of *D. dubosii* depended specifically on the concentration of Trp but not on other amino acids (Fig. 4I, J). Based on these experimental results, we have paid more attention to the first hypothesis. These results suggest that Trp supplementation results in an increase in *D. dubosii*, which may explain the enhanced survival with Trp supplementation in tumor-bearing mice.

### *D. dubosii* replicates the Trp-induced enhanced antitumor response against brain tumor

To verify the role of *D. dubosii* in Trp-mediated enhanced survival in mice with brain tumor, we colonized GF mice with this strain (Fig. 5A). This single colonization with *D. dubosii* increased survival against brain tumor (Fig. 5B). In addition, tumor burden was also reduced in colonized GF mice (Fig. 5C). Remarkably, these effects of *D. dubosii* colonization were comparable to those observed with Trp treatment (Fig. 2F, G and Fig. 5B, C). This effect depended on colonization with live bacteria because heat-killed *D. dubosii* didn’t replicate the survival improvement (Additional file 1: Fig. S8A).

**Fig. 5.**
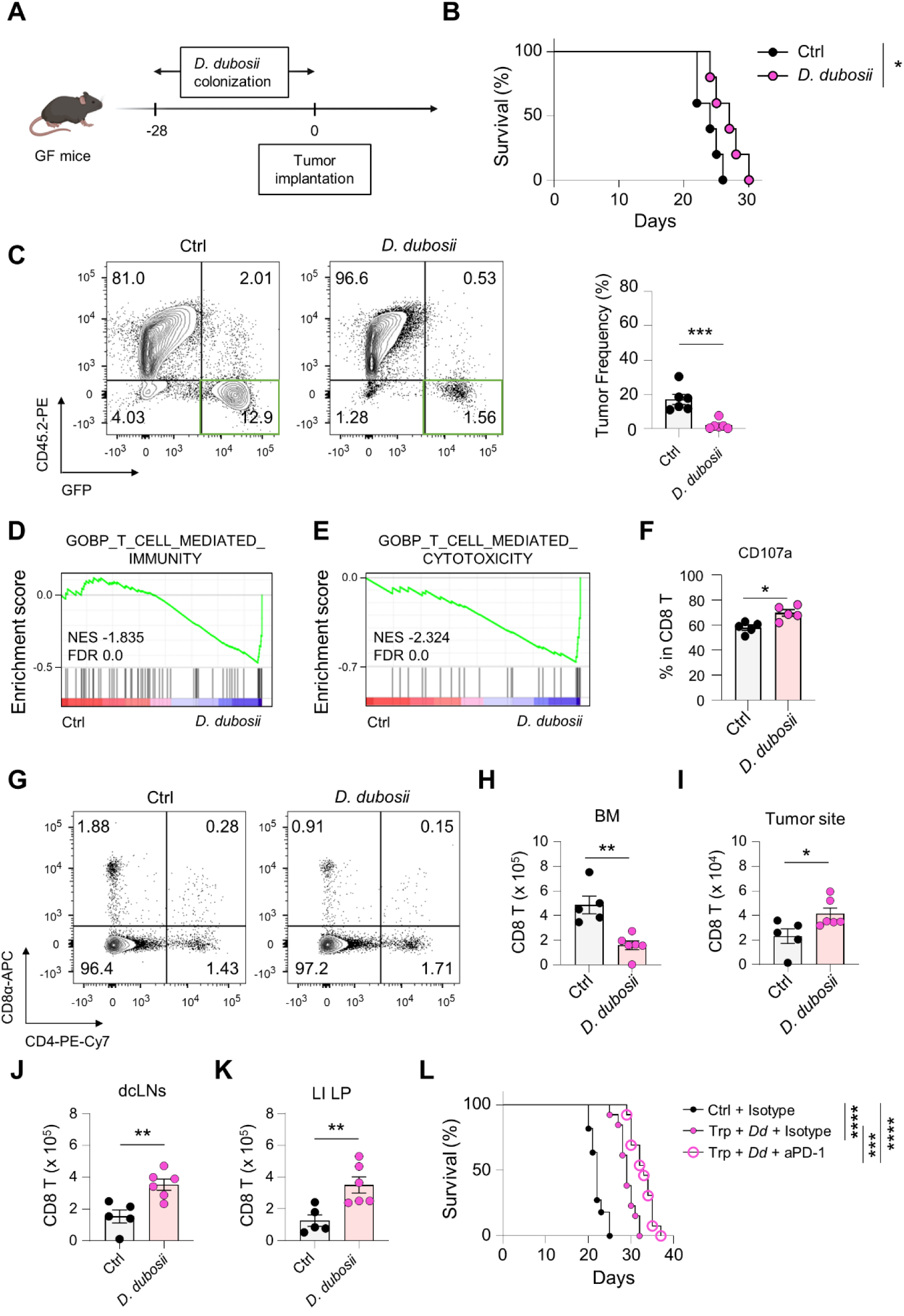
*D. dubosii* mimicked the Trp-mediated enhancement of the antitumor response against brain tumors. **A** Four weeks after the colonization of GF mice with *D. dubosii*, GL261-GFP tumor cells were transplanted into the right hemisphere of the mice, and **B** Survival was measured for the colonized and the control groups (n *=* 5 for each group). **C** On D20, tumor burden was measured by flow cytometry. **D**, **E** On D20 after tumor transplantation, immune cells were sorted from the brains of tumor-bearing colonized mice (*D. dubosii*) and the control (Ctrl) mice. GSEA analysis used gene set (**D**) GOBP_T_CELL _MEDIATED_IMMUNITY and (**E**) GOBP_T_CELL_MEDIATED_CYTOTOXICITY in a tumor-infiltrating CD8 T-cell subset comparing the colonized mice (*D. dubosii*) and the control (Ctrl) mice. The transcriptome was analyzed using scRNA-seq. **F** CD107a expression was determined by flow cytometry on D20. **G**-**K** On D20 after tumor transplantation for the colonized (*D. dubosii*) and the control (Ctrl) groups, CD8 and CD4 T cells were quantified by flow cytometry in bone marrow (**G**, **H**). Similarly, CD8 T cells were measured in the brain (**I**), deep cervical lymph nodes (**J**), and colon lamina propria (**K**). **L** Survival of tumor-bearing mice after *D. dubosii* colonization (*Dd*) with or without anti-PD-1 (aPD-1) treatment was determined with supplemental Trp provided to improve colonization. Ctrl, n *=* 11; Trp + *Dd* +isotype and Trp + *Dd* + aPD-1, n *=* 13. Statistical significance was calculated using a two-tailed unpaired Student’s *t*-test in (**C**, **F**, **H**-**K**) or a log-rank test in (**B**, **L**). Data are presented as mean ± s.e.m. *\*P <* 0.05, \**\*P <* 0.01, \*\**\*P <* 0.001, \*\*\**\*P <* 0.0001.

Next, transcriptomic analysis showed an increase in CTL activity in mice colonized with *D. dubosii*, as evidenced by the upregulation of CD107a, a key marker of cell-mediated cytotoxicity (Fig. 5D–F and Additional file 1: Fig. S8B–C). However, the expression of T cell exhaustion markers was similar in *D. dubosii*–colonized and control mice (Additional file 1: Fig. S8D–F). T cell sequestration was also affected by colonization with *D. dubosii* (Fig. 5G–K), and anti-PD-1 therapy in mice colonized with *D. dubosii* further increased survival (Fig. 5L). In GF mice, sole anti-PD-1 treatment didn’t show the survival improvement [21]. Thus, the increase in abundance of *D. dubosii* in the gut affected T cell circulation and enhanced the T cell mediated antitumor immune response, leading to improved survival in mice with brain tumor.

## Discussion

Here, we characterized the distinctive changes in the gut microbiota that occur during the progression of brain tumors. To modulate gut microbiota, we fed Trp to mice with brain tumors and observed an increase in their survival, which was dependent on the gut microbiota. This increased survival rate was associated with alterations in T cell circulation and increased T cell infiltration. Further, feeding mice with Trp increased the efficacy of anti-PD-1 antibody immunotherapy. We identified the bacterial strain *D. dubosii* that correlated best with the observed changes in the tumor-bearing mice fed Trp, and showed that providing *D. dubosii* to GF mice replicated the phenotype observed by feeding mice with Trp.

Although numerous studies have investigated the gut-brain connection [28–31], establishing causal effects, especially in cancer, remains challenging due to the distance between these organs. Despite direct implications of the microbiota in various cancers [32], the mechanisms underlying their effects remain elusive. Our study suggests that translocation events involving gut microbiota are improbable, as our treatment did not alter the intestinal permeability barrier. Instead, we propose that observed effects on brain tumors result from changes in the circulation of specific immune cells, particularly CD8 T cells, challenging the conventional involvement of the nervous and endocrine systems in the gut-brain axis.

Immunotherapy has limited efficacy in cold tumors such as brain tumors, making them difficult to treat [33]. This is partly due to the fact that in brain tumors, T cells are sequestered in the bone marrow, thereby reducing T cell circulation and the efficacy of antibodies that target immune checkpoint molecules [8]. Here, we show that modulation of gut microbiota can improve the efficacy of immunotherapy. Although further research is needed to identify specific targets, this study provides important insights into the enigma of brain tumors and potential approaches to address the challenges of treating them. As this sequestration is also observed in human GBM patients [8], our results could be translated into human cases.

In our study, we focused on the decrease in gut Trp levels during brain tumor progression and its association with changes in gut microbiota. To address this, we used Trp supplementation, despite the unclear cause of this reduction. Quantitative changes in amino acids can be diagnostic markers in certain diseases, including brain tumors. However, the poor prognosis of GBM highlights the need for more effective treatments. Understanding the cause of Trp reduction could lead to new avenues. While our findings suggest that supplemented Trp affects fecal rather than brain Trp levels, further investigation into intracellular and extracellular Trp levels in tumor cells is needed. Tumor cells often metabolize Trp, and exploring the effects of other amino acids could significantly impact brain tumor research.

While our research has contributed significantly to understanding the gut microbiota’s role in brain tumors, further studies are necessary. 1) Little is known in humans about the bacterial strains that we identified here in mice [34, 35]. However, as T cell sequestration occurs in human GBM patients, further research using these bacterial strains in humans is warranted. 2) Although we did not study changes in serotonin levels here, supplemental Trp could induce neurological change via serotonin [36]. 3) We identified 19 bacterial strains as candidates to explain the Trp-mediated effect on brain tumor; however, we were unable to classify most of them at the species level. It is possible that some of these uncharacterized strains affect the host defense against brain tumors by other mechanisms.

## Conclusion

Our findings facilitate the expansion of the gut-brain axis to encompass tumors. Moreover, instead of predominantly addressing established host-host interactions, we examine the interplay between gut microbiota and the host in this axis. Additionally, we present a strategy to enhance the efficacy of third-generation cancer therapies in brain tumors, exemplifying cold tumors. Moreover, the investigated dietary intervention and the utilization of *D. dubosii* in this study offer potential considerations for the treatment of brain tumor patients or lifestyle modifications.

## Methods

### Animals

Male specific pathogen-free (SPF) C57BL/6J mice, aged between 5 to 8 weeks, were obtained from Dae Han Bio Link Co. Ltd (Eumseong, Korea). Germ-free (GF) mice were acquired from POSTECH Biotech Center (Pohang, Korea). LSL-EGFRvIII mice were provided by Dr. Jeong Ho Lee (KAIST, Korea). Athymic nude mice (Koat:Athymic NCr^-nu/nu^) were purchased from Koatech (Pyeongtaek, Korea). All mice were housed in KAIST SPF facility under a 12-hour light/dark cycle at 18–24°C and 30–70% humidity. Experimental procedures adhered to KAIST IACUC guidelines (KA2020-126, KA2021-118, and KA2021-119). Experimental mice received 0.2% Trp-supplemented drinking water starting three weeks prior to tumor inoculation and throughout the study.

### Bacteria and cell lines

*Duncaniella dubosii* (*D. dubosii*; KCTC15734) was obtained from the Korean Collection for Type Cultures (KCTC, Jeongeup, Korea) [37]. This was cultured in brain heart infusion (BHI; HiMedia, Kennett Square, PA, USA) medium supplemented with hemin (TCI, Tokyo, Japan) and vitamin K (Alfa Aesar, Ward Hill, MA, USA) (BHIS), using an anaerobic system comprising an anaerobic jar, anaerogen, and indicator (Oxoid, Basingstoke, UK). Bacterial colony-forming units (CFUs) or cells were counted on BHI agar plates or using a hemocytometer (Hausser Scientific, Horsham, PA, USA), respectively. For storage, bacteria were preserved in 30% glycerol (Samchun Chemicals, Seoul, Korea) in a BHIS medium and stored at -80℃.

For growth comparison, 5 x 10^8^ cells/mL of *D. dubosii* were cultured in BHIS medium with or without 0.1, 1, and 10 mM of Trp for 5 days using anaerobic system. In the second experiment, at 10 mM concentration, the influence of amino acids was compared.

GL261 mouse glioma cell lines and green fluorescent protein (GFP)-tagged GL261 (GL261-GFP) were provided by Dr. Injune Kim (KAIST) [38]. These cells were passaged with trypsin/EDTA (Welgene; Gyeongsan, Korea) and cultured in Dulbecco’s modified Eagle’s medium (DMEM; Corning, Corning, NY, USA) supplemented with 10% fetal bovine serum (FBS) and 1% penicillin-streptomycin (Welgene).

### Brain tumor animal models

For the gene-editing GBM tumor model, murine GBM was induced spontaneously using a Cre-loxP and Crispr-Cas9 system [39]. Vectors for GBM induction, provided by Dr. Jeong Ho Lee (KAIST), contained single-guide RNA (sgRNA) for two tumor suppressor genes (*TP53* and *PTEN*), *Cas9*, and *Cre* recombinase. These vectors were transferred into P1-P2 pup’s ventricle through electroporation.

In the intracranial implantation model [39], 8-week-old male mice received injections of either 1 × 10^5^ GL261 or GL261-GFP cells. Mice were anesthetized using isoflurane, positioned in a stereotaxic apparatus (Stoelting Co., Wood Dale, IL, USA), and a hole was drilled in the skull. Tumor cells were injected into the right frontal cortex using a Hamilton syringe (Hamilton Company, Reno, NV, USA). For sham injection, DPBS was injected instead. In this model, the stereotaxic apparatus was utilized within a biosafety level 2 biocabinet (CHC LAB, Daejeon, Korea) to prevent exposure to undesired microbes.

### Bacterial Colonization

For bacterial colonization in GF mice, we administered 1 × 10^8^ CFU of bacteria in 100 μL of 1.5% sodium bicarbonate *per os* using a Zonde needle (20G, 0.9 × 50 mm; Jeung do Bio & Plant Co. Ltd, Seoul, Korea) for four weeks before tumor implantation using a schedule of five consecutive days of administration followed by a 2-day break.

For fecal microbiota transplantation (FMT), we collected and passed 50 mg of feces through a 70 μm strainer (SPL, Korea), centrifuged the filtrate, suspended the stool pellet in 500 μL of DPBS, and administered 100 μL of the suspension to each recipient *per os*. FMT was also done for four weeks before tumor injection, as described above.

### Gut permeability test

To assess the gut integrity, we administered 500 mg/kg of 4 kDa fluorescein isothiocyanate– dextran (FITC-Dextran; Sigma-Aldrich, St. Louis, MO, USA) *per os* to mice after a 4-hour fast. After 4 h, blood was collected through submandibular vein. Plasma was prepared after centrifugation (5000 rpm for 10 mins) from blood collected in heparin-coated anti-coagulation tubes (BD Biosciences, Bergen County, NJ, USA), and fluorescence of the diluted plasma samples was measured using a SPECTROSTAR Nano Microplate Reader (BMG Labtech, Ortenberg, Germany) with a 485 nm excitation wavelength and a 530 nm emission wavelength.

### Microbiome analysis

Fecal samples were collected from mice, and bacterial DNA was extracted using the QIAamp DNA Stool Mini Kit (QIAGEN). The V3-V4 regions of the 16S rRNA gene were amplified using 341F and 805R primers. Sequencing was performed on the Illumina Miseq system. Primer sequences was following: 341F (5’-AATGATACGGCGACCACCGAGATC TACAC-XXXXXXXX-TCGTCGGCAGCGTCAGATGTGTATAAGAGACAG-CCTACGGGNGGCWGCAG-3’) and 805R (5’-CAAGCAGAAGACGGCATACGAGAT-XXXXXXXXGTCTCGTGGGCTCGG-AGATGTGTATAAGAGACAG-GACTACHVGGGTATCTAATCC-3’).

For data processing, raw read quality was assessed using Trimmomatic, with low-quality reads (<Q25) removed. Paired-end data were merged, and unique reads were extracted and clustered using VSEARCH. The EzBioCloud 16S rRNA database served for taxonomic assignment [40], with chimeric reads filtered out by UCHIME algorithm. Alpha diversity indices (ACE, Chao1, Jackknife, Shannon, NPShannon, Simpson, Phylogenetic diversity) and generalized UniFrac algorithms for beta diversity were calculated. Functional analysis was conducted using PICRUSt, identifying functional and taxonomical microbial markers. The LEfSe test and Kruskal-Wallis H Test were utilized for heatmap analysis.

### Tissue homogenization

Sliced brain tissue was filtered through a 70-μm cell strainer in 1 mL of DPBS. After centrifugation (5000 rpm for 5 mins at 4℃), the supernatant was utilized. Feces (50 mg) were mixed with 450 μL of DPBS, vigorously vortexed for 1 min, and then incubated on ice for 5 mins, followed by another 1-minute vortex. The supernatant was collected after 2-minute ice incubation through centrifugation (10000 x g, 4°C). Blood from mandibular veins was incubated at 37°C for 1 h, then centrifuged at 10,000 rpm for 5 mins to isolate serum.

### Amino acid measurements

To measure amino acids, supernatant from homogenized feces was used for quantification using a 15 Tesla Fourier transform ion cyclotron resonance mass spectrometry (15T FT-ICR MS) system (Bruker Daltonics, Billerica, MA, USA). We injected 2 μL of standard amino acids (Sigma-Aldrich) at concentrations of 1, 2, 10, 20, 50, and 100 pmol and read them at a flow rate of 0.5 mL/min at a wavelength of 260 nm.

The amount of Trp in homogenized samples was measured using an enzyme-linked immunosorbent assay (ELISA) following the manufacturer’s instructions (Novus, Chesterfield, MO, USA). Optical density was measured at 450 nm (Bio-Rad) using SpectraMax microplate reader (Molecular Devices, San Jose, CA, USA), and the concentration of Trp was calculated using standard curves.

### CD8 T cell in vivo depletion

For CD8α^+^ cell depletion, 150 μg of anti-CD8α (2.43; BioXcell, Lebanon, NH, USA) in 200 μL of DPBS was injected into mice intraperitoneally. The isotype control was monoclonal Rat IgG2b (LTF-2; BioXcell). Mice were injected with antibodies 1 and 2 days before tumor implantation and every 7 days after tumor implantation.

### Single cell preparation

For flow cytometry and transcriptomic analysis, single-cell suspensions were prepared from brain tissue [39], lymph nodes [39], bone marrow tissue, and colon. Tissues were harvested from mice following euthanasia via a CO_2_ chamber and cardiac perfusion with 50 mL of cold DPBS.

Femurs and tibias were isolated from the hind legs of euthanized mice to obtain bone marrow cells. The bone marrow cells were harvested by flushing the DMEM medium supplemented with 1% FBS through the bones after cutting the ends. Red blood cell lysis was performed using ACK lysis buffer.

Isolated colon tissues were prepared by removing fats and Peyer’s patches. The tissues were washed thrice with 2% FBS in RPMI 1640 medium (Corning). They were then cut into rectangular sections. Digestion was performed using the GentleMACS Dissociator with the Lamina Propria Dissociation Kit (Miltenyi Biotec). Immune cells were isolated using a Percoll gradient, and RBCs were lysed with ACK buffer.

### Flow cytometry

Single-cell suspensions were treated with anti-CD16/32 antibody (2.4G2) to block Fc receptors and stained with the antibodies for: anti-CD45.2, anti-CD3ε, anti-CD4, anti-CD8α, anti-PD-1, anti-TIM-3. Dead cells were separated from live cells using Fixable Viability Dye eFluor 450 (Thermo Fisher Scientific, Waltham, MA, USA) or propidium iodide (Invitrogen, Carlsbad, CA, USA).

To activate Tumor-infiltrating lymphocytes (TILs) [26], phorbol-myristate acetate (PMA; Sigma-Aldrich) and ionomycin (Sigma-Aldrich) were applied for 4 h at 37°C with GolgiStop (BD Biosciences) and GolgiPlug (BD Biosciences). Intracellular staining was performed using Foxp3 Fix/Perm Buffer Set (Biolegend) after surface staining. Cytokine staining included anti-IFN-γ and anti-TNF-α antibodies. For cytotoxicity comparison, anti-CD107a staining was conducted. Data acquisition was done using an LSRFortessa flow cytometer (BD Biosciences) and analyzed using FlowJo software (v.10.5.3; Treestar, Ashland, OR, USA). Detailed information for antibodies is in Table S1 (Additional file 1).

### Single-cell transcriptome analysis

For transcriptomic analysis, viable CD45.2^+^ cells negative for 7-amino actinomycin D (7-AAD; BD Biosciences) were sorted from brain tumors supplemented with Trp using fluorescence-activated cell sorting ARIA II (BD Biosciences). Similarly, cells were sorted from tumor-bearing mice colonized with *D. dubosii* using FACS ARIA Fusion cell sorters (BD Biosciences). Single-cell libraries were generated using the 10× Chromium Single Cell 3’ Gene Expression kit, with 10,000 cells from each sample sequenced on the HiSeqXten (Illumina). Generation of Seurat objects using sequencing output files was conducted using Seurat R package [26]. Here, we analyzed 8,107 control and 4,788 Trp-treated cells, and 8768 control and 8913 *D. dubosii*– colonized cells. Differential gene expression was identified and visualized, and gene set enrichment analysis (GSEA) was conducted using software [41] (GSEA v.4.1.0; Broad Institute, Cambridge, MA, USA).

### Immunotherapy

For immunotherapy using an anti-PD-1 antibody, we injected 200 μg of anti-PD-1 blocking antibody (RMP1-14; BioXcell) into mice intraperitoneally on days 10, 13, and 16 after tumor inoculation. Rat IgG2a isotype antibody (2A3; BioXcell) was used as control and injected using the same schedule.

### Statistical analysis

Data are expressed as means ± standard error mean (s.e.m.). Group differences were assessed using an unpaired, two-tailed Student’s *t*-test. Survival disparities were evaluated using the log-rank test, and correlations were analyzed using Pearson’s correlation test. Statistical analyses were performed using Prism Software (v.9.0.5; GraphPad; San Diego, CA, USA) and R software (R Foundation for Statistical Computing). A significance level of P < 0.05 was applied.

## Availability of data and materials

Single-cell RNA sequencing data were deposited in the Gene Expression Omnibus (GEO) at the National Center for Biotechnology Information (NCBI) with accession code GSE256108.

## Acknowledgements

We thank all the members of the Host Defense lab for helpful discussions and Ji Ye Kim at the BioMedical Research Center for technical assistance.

## Author contributions

H.C.K, H.-J.K, J.L, W.H.P, S.H.P, B.H.K and Y.K. conducted the experiments. H.C.K and H.K.L designed the experiments, analyzed the data, and wrote the manuscript. H.K.L. supervised the study.

## Ethics declarations

### Ethics approval and consent to participate

All procedures treated with mice were approved by the Institutional Animal Care and Use Committee of KAIST. Experimental procedures adhered to KAIST IACUC guidelines (KA2020-126, KA2021-118, and KA2021-119).

## Consent for publication

Not applicable.

## Additional file 1

Filename: Kim-et-al-Supplementary Additional-File-0403-2024final Contents: Figure S1-8 and Table S1

Title of Data:

**Figure S1.** In two GBM animal models, gut microbiota alpha diversity remained similar as tumors progressed.

**Figure S2.** As the brain tumor progresses, the amount of tryptophan decreased.

**Figure S3.** Transcriptomic analysis reveals a stronger T-cell response in Trp-treated mice with similar CTLs.

**Figure S4.** Gating strategy for the analysis of T cell effector or exhaustion molecule expression.

**Figure S5.** At early time points, T cells infiltrate the TME in brain tumors.

**Figure S6.** Trp supplementation enhances the circulation of CD8 T cells.

**Figure S7.** The Trp-mediated effect depends on the gut microbiota.

**Figure S8.** Trp-mediated changes in CTLs are observed after colonization of GF mice with *living D. dubosii*.

**Table S1.** Lists for flow cytometric antibody

Description of Data:

**Figure S1.** In two GBM animal models, gut microbiota alpha diversity remained similar as tumors progressed.

**A** We used a Cre-loxP and Crispr-Cas9 system to generate genetically engineered spontaneous mouse GBM model. The vectors to create the GBM mouse strain contained single-guide RNA (sgRNA) for two tumor suppressor genes, *TP53* and *PTEN*, *Cas9*, and *Cre* recombinase. After we injected the vectors into the right lateral ventricle of 1–2-day old (P1-P2) EGFRvIII mice pups, we applied an electrical pulse for electroporation. **B** For the orthotopic implantation model, mice were anesthetized with diluted isoflurane (1:1 with oxygen) and held in a stereotaxic apparatus. We drilled a hole in the right hemisphere of the skull at a position 2 mm lateral to and 2 mm posterior from the bregma. We injected 1 × 10^5^ tumor cells into the right frontal cortex to a depth of 3 mm depth using a syringe over 5 mins. During this intracranial injection, mice were fixed on a stereotaxic instrument. **C**, **D** At the indicated time points, fecal samples were collected and analyzed by 16S rRNA sequencing. **E**-**H** We compared the alpha diversity of the gut microbiota for both models as the tumor progressed, measuring both species richness (**E**, **F**) and diversity (**G**, **H**). Genetically engineering model (GEM model): (**E**, **G**), Orthotopic implantation model: (**F**, **H**). For statistical analysis, a two-tailed, unpaired Student’s *t*-test was used. Data are presented as mean ± s.e.m. \*\**P* < 0.01, ns: not significant.

**Figure S2.** As the brain tumor progresses, the amount of tryptophan decreased.

**A**-**D** Mice received implantation of GL261 cells into their right hemispheres. Blood and fecal samples were collected either before tumor injection or 10 or 20 days after (**A**). Serum (**B**) and fecal tryptophan (**C**) were measured by ELISA in samples collected at the time points indicated in (**D**). The correlation between the amount of serum and fecal tryptophan was calculated with linear regression. **E**-**L** Using the PICRUSt algorithm, the relative abundance of bacterial modules (**E**-**H**) and pathways (**I**-**L**) were compared. **E**-**H** Modules showing differences at a significant level (*P* < 0.05) were selected. Among these modules, the top 10 exhibiting the most significant increase were highlighted in red, while the top 10 showing the most significant decrease were marked in blue. These comparison was conducted in two brain tumor animal model: GEM model (**E**, **G**, **I**, **J**), orthotopic implantation model (**F**, **H**, **K**, **L**). Statistical significance was calculated using a two-tailed, unpaired Student’s *t*-test. Data are presented as mean ± s.e.m. \**P* < 0.05, \*\**P* < 0.01, \*\*\**P* < 0.001, \*\*\*\**P* < 0.0001, ns: not significant.

**Figure S3.** Transcriptomic analysis reveals a stronger T-cell response in Trp-treated mice with similar CTLs.

**A**, **B** The transcriptomes of sorted CD45.2^+^ immune cells from brain tissue taken on D20 from DW vs. Trp mice were analyzed using single-cell RNA sequencing. Immune cells were clustered by their marker genes, and the expression of effector molecules (**E**) and exhaustion molecules (**H**) on CTLs was compared between the two groups. (**C**, **D**, **F**, **G**) The expression of related proteins was measured by flow cytometry. **I**-**K** From tumor-bearing mice under Trp treatment, the number of brain cells was compared with control mice’ (**I**). CD8 T cells from the brain were quantified on D20 using flow cytometry, including the number (**J**) and frequency among CD3ε^+^ cells (**K**) (n = 5 per group). We used a two-tailed, unpaired Student’s *t*-test for statistical comparison. Data are presented as mean ± s.e.m. \**\*P <* 0.01, \*\*\*\**P* < 0.0001, ns: not significant.

**Figure S4.** Gating strategy for the analysis of T cell effector or exhaustion molecule expression.

The gating was performed in the order depicted in the figure. The red line represents the gating flow of expression for effector molecules, and the blue line represents that of expression for exhaustion molecules.

**Figure S5.** At early time points, T cells infiltrate the TME in brain tumors.

For temporal analysis, control samples and TME samples (d10 and d20) prepared D10 or D20s after tumor inoculation were integrated. The infiltration of CD8 T cells (**A**) and CD4 T cells (**B**) was assessed by the expression of *Cd8α* and *Cd4*, respectively.

**Figure S6.** Trp supplementation enhances the circulation of CD8 T cells.

**A** Gating strategy for T-cell gating. **B**, **C** D10 (n = 5) or D20 (n = 4) after tumor implantation, bone marrow cells from tumor-bearing mice were analyzed using flow cytometry using bone marrow of healthy mice (n = 3) as the control. **D**-**G** On D20, T cells were quantified in bone marrow (**D**), tumor site (**E**), deep cervical lymph nodes (**F**), and large intestine lamina propria (**G**) by flow cytometry from control (DW, n = 4) and Trp-treated tumor-bearing mice (n = 4). We used a two-tailed, unpaired Student’s *t*-test for statistical significance comparison. Data are presented as mean ± s.e.m. *\*P <* 0.05, \**\*P <* 0.01, ns: not significant.

**Figure S7.** The Trp-mediated effect depends on the gut microbiota.

**A**, **B** The expression of *Eif2ak4* (GCN2) was compared in the control (DW) and Trp-treated group (Trp) for total tumor-infiltrating lymphocytes (TILs) (**A**) and CTLs (**B**). **C**, **D** GSEA analysis for HALLMARK_APOPTOSIS in T cells (**C**) and B cells (**D**). **E** Survival was determined for GF mice colonized with fecal microbiota from control (DW) or Trp-treated tumor-bearing mice (Trp) for 4 weeks before tumor cells were implanted into their brains. **F** To measure gut barrier integrity on D0 and D20 after tumor implantation, mice that had fasted for 4 hours were treated with 4 kDa dextran-FITC *per os,* and 4 hours later, fluorescence was measured in the plasma. For statistical analysis, a log-rank test was used in (**E**), and a two-tailed unpaired Student’s *t*-test was used in (**F**). Data are presented as mean ± s.e.m. *\*P <* 0.05, ns: not significant.

**Figure S8.** Trp-mediated changes in CTLs are observed after colonization of GF mice with *living D. dubosii*.

**A** Survival of GF mice colonized with heat-killed *D. dubosii per os* for 8 weeks, starting from 4 weeks before tumor inoculation, was measured after tumor implantation. **B**, **C** Transcriptomic analysis of tumor infiltrating leukocytes from GF tumor-bearing mice colonized or not with *D. dubosii*. **D**-**F** The expression of exhaustion molecules was measured using flow cytometry (**D**, **E**) and single-cell RNA sequencing (**F**). Both analysis were conducted on D20 (n =5 for each group). For statistical analysis, a log-rank test was used in (**A**), a two-tailed unpaired Student’s *t*-test was used in (**D**-**F**). Data are presented as mean ± s.e.m. ns: not significant.

**Table S1.** Lists for flow cytometric antibody

