## Supplementary Figure S1 for "Gut microbiota dysbiosis induced by brain tumor modulates the efficacy of immunotherapy"

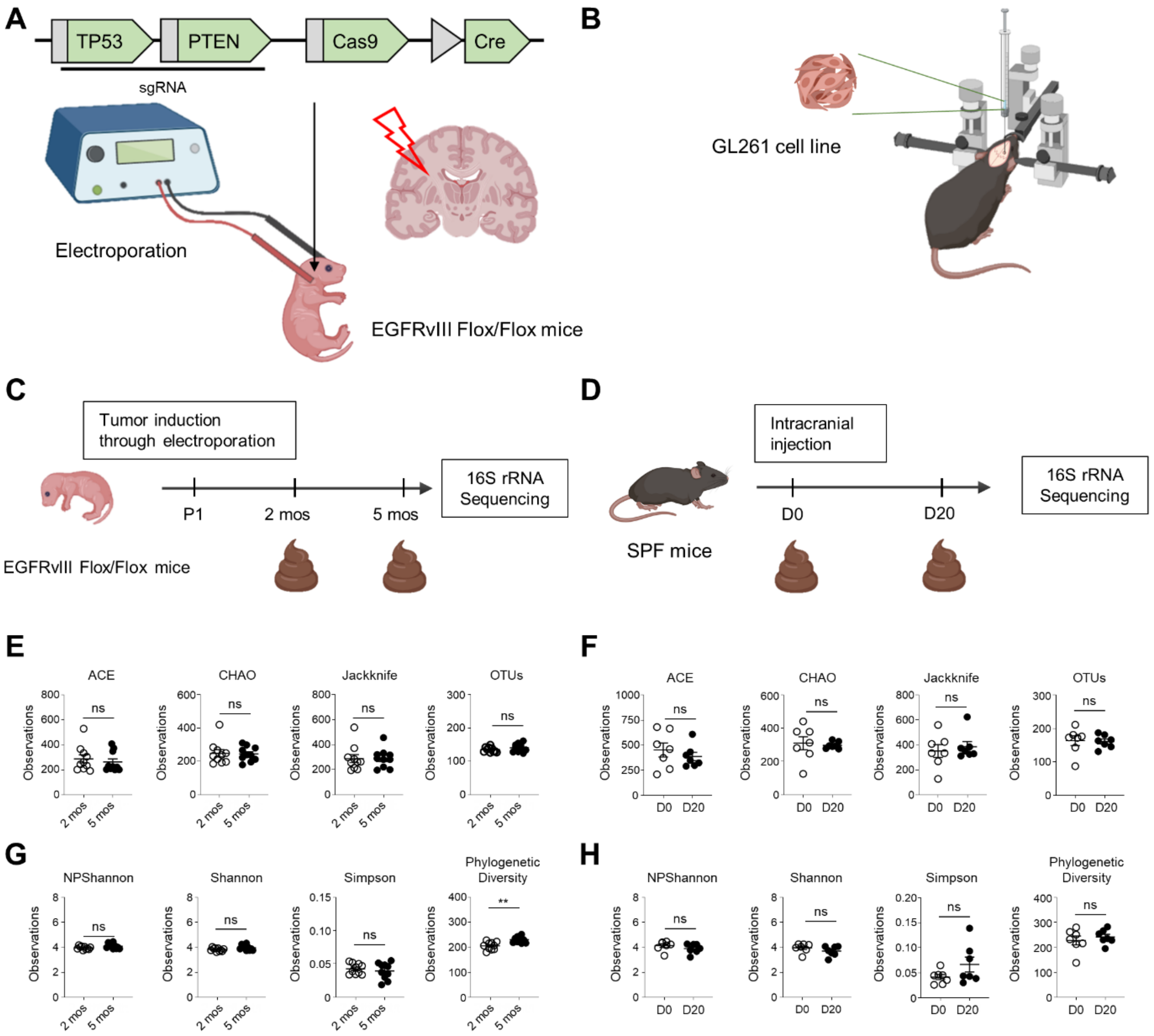


**Figure S1.** In two GBM animal models, gut microbiota alpha diversity remained similar as tumors progressed.

**A** We used a Cre-loxP and Crispr-Cas9 system to generate genetically engineered spontaneous mouse GBM model. The vectors to create the GBM mouse strain contained single-guide RNA (sgRNA) for two tumor suppressor genes, *TP53* and *PTEN*, *Cas9*, and *Cre* recombinase. After we injected the vectors into the right lateral ventricle of 1–2-day old (P1-P2) EGFRvIII mice pups, we applied an electrical pulse for electroporation. **B** For the orthotopic implantation model, mice were anesthetized with diluted isoflurane (1:1 with oxygen) and held in a stereotaxic apparatus. We drilled a hole in the right hemisphere of the skull at a position 2 mm lateral to and 2 mm posterior from the bregma. We injected 1 × $10^5$ tumor cells into the right frontal cortex to a depth of 3 mm depth using a syringe over 5 mins. During this intracranial injection, mice were fixed on a stereotaxic instrument. **C**, **D** At the indicated time points, fecal samples were collected and analyzed by 16S rRNA sequencing. **E**-**H** We compared the alpha diversity of the gut microbiota for both models as the tumor progressed, measuring both species richness (**E**, **F**) and diversity (**G**, **H**). Genetically engineering model (GEM model): (**E**, **G**), Orthotopic implantation model: (**F**, **H**). For statistical analysis, a two-tailed, unpaired Student's *t*-test was used. Data are presented as mean ± s.e.m. **$P < 0.01$, ns: not significant.
