## Supplementary Figure S2 for "Gut microbiota dysbiosis induced by brain tumor modulates the efficacy of immunotherapy"

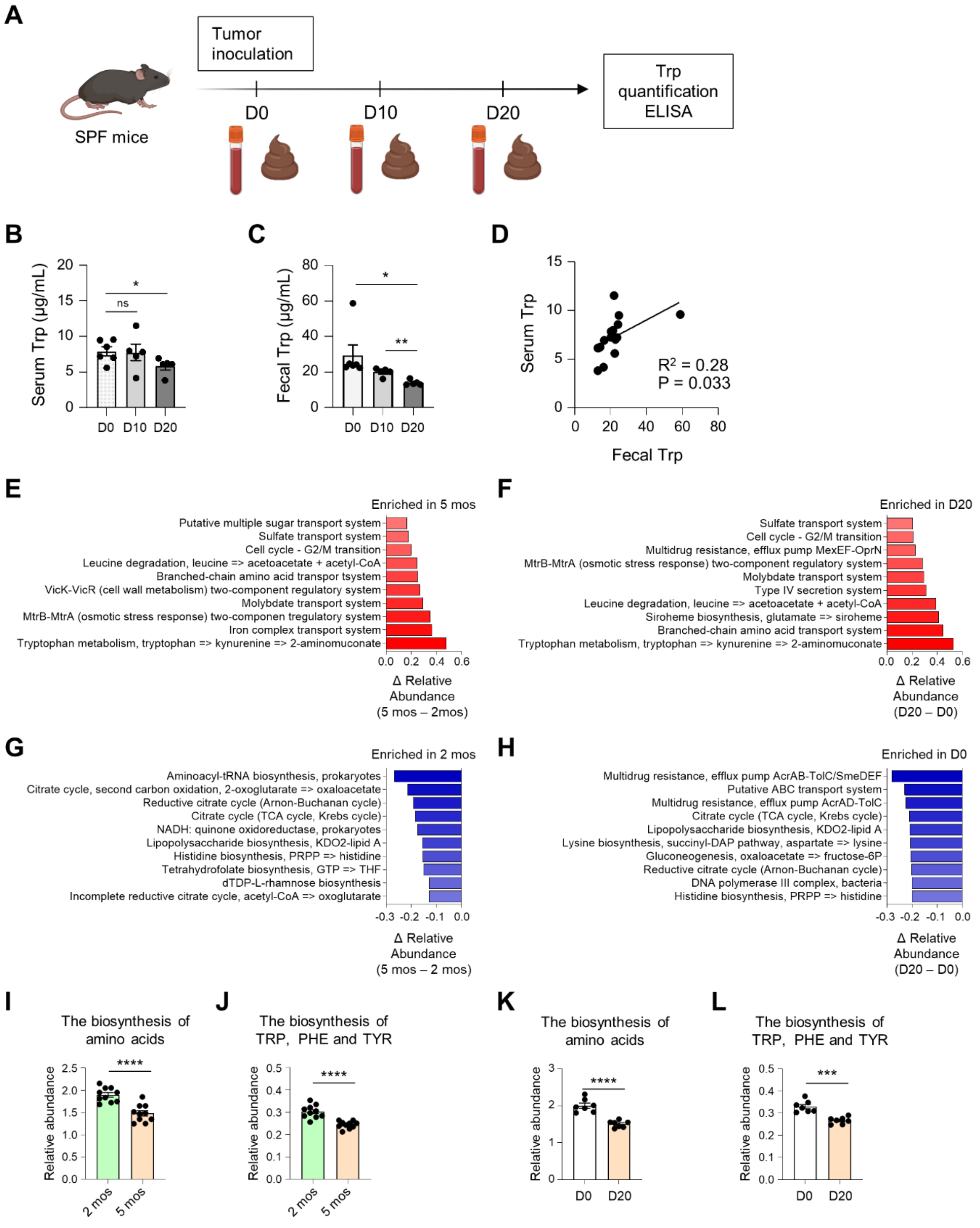


**Figure S2.** As the brain tumor progresses, the amount of tryptophan decreased.

**A**-**D** Mice received implantation of GL261 cells into their right hemispheres. Blood and fecal samples were collected either before tumor injection or 10 or 20 days after (**A**). Serum (**B**) and fecal tryptophan (**C**) were measured by ELISA in samples collected at the time points indicated in (**D**). The correlation between the amount of serum and fecal tryptophan was calculated with linear regression. **E**-**L** Using the PICRUSt algorithm, the relative abundance of bacterial modules (**E**-**H**) and pathways (**I**-**L**) were compared. **E**-**H** Modules showing differences at a significant level ($P < 0.05$) were selected. Among these modules, the top 10 exhibiting the most significant increase were highlighted in red, while the top 10 showing the most significant decrease were marked in blue. These comparison was conducted in two brain tumor animal model: GEM model (**E**, **G**, **I**, **J**), orthotopic implantation model (**F**, **H**, **K**, **L**). Statistical significance was calculated using a two-tailed, unpaired Student's *t*-test. Data are presented as mean ± s.e.m. *$P < 0.05$, **$P < 0.01$, ***$P < 0.001$, ****$P < 0.0001$, ns: not significant.
