## Supplementary Figure S3 for "Gut microbiota dysbiosis induced by brain tumor modulates the efficacy of immunotherapy"

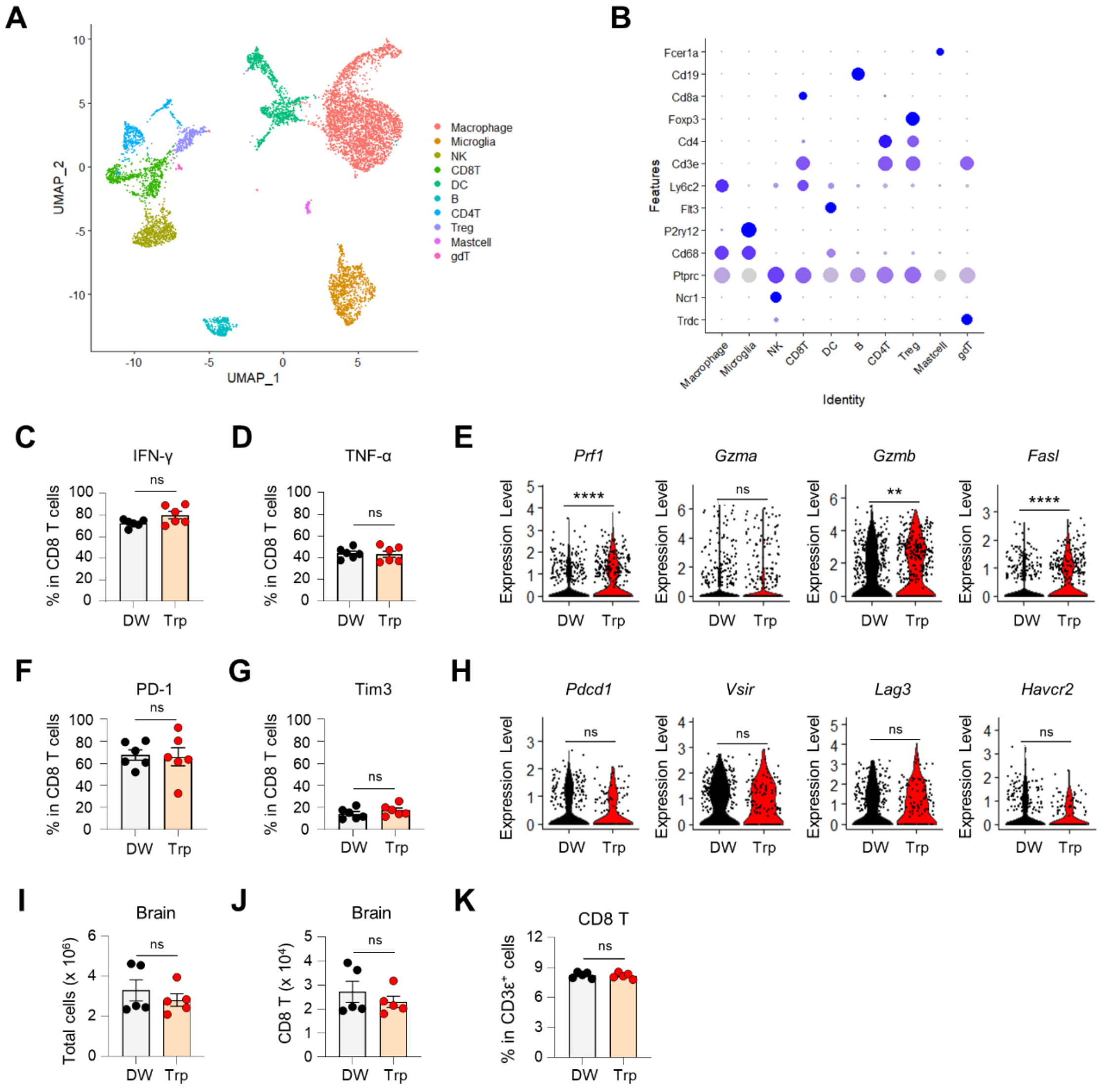


**Figure S3.** Transcriptomic analysis reveals a stronger T-cell response in Trp-treated mice with similar CTLs.

**A**, **B** The transcriptomes of sorted $CD45.2^+$ immune cells from brain tissue taken on D20 from DW vs. Trp mice were analyzed using single-cell RNA sequencing. Immune cells were clustered by their marker genes, and the expression of effector molecules (**E**) and exhaustion molecules (**H**) on CTLs was compared between the two groups. (**C**, **D**, **F**, **G**) The expression of related proteins was measured by flow cytometry. **I-K** From tumor-bearing mice under Trp treatment, the number of brain cells was compared with control mice' (**I**). CD8 T cells from the brain were quantified on D20 using flow cytometry, including the number (**J**) and frequency among CD3ε$^+$ cells (**K**) (n = 5 per group). We used a two-tailed, unpaired Student's *t*-test for statistical comparison. Data are presented as mean ± s.e.m. $^{**}P < 0.01$, $^{****}P < 0.0001$, ns: not significant.
