## Supplementary Figure S4 for "Gut microbiota dysbiosis induced by brain tumor modulates the efficacy of immunotherapy"

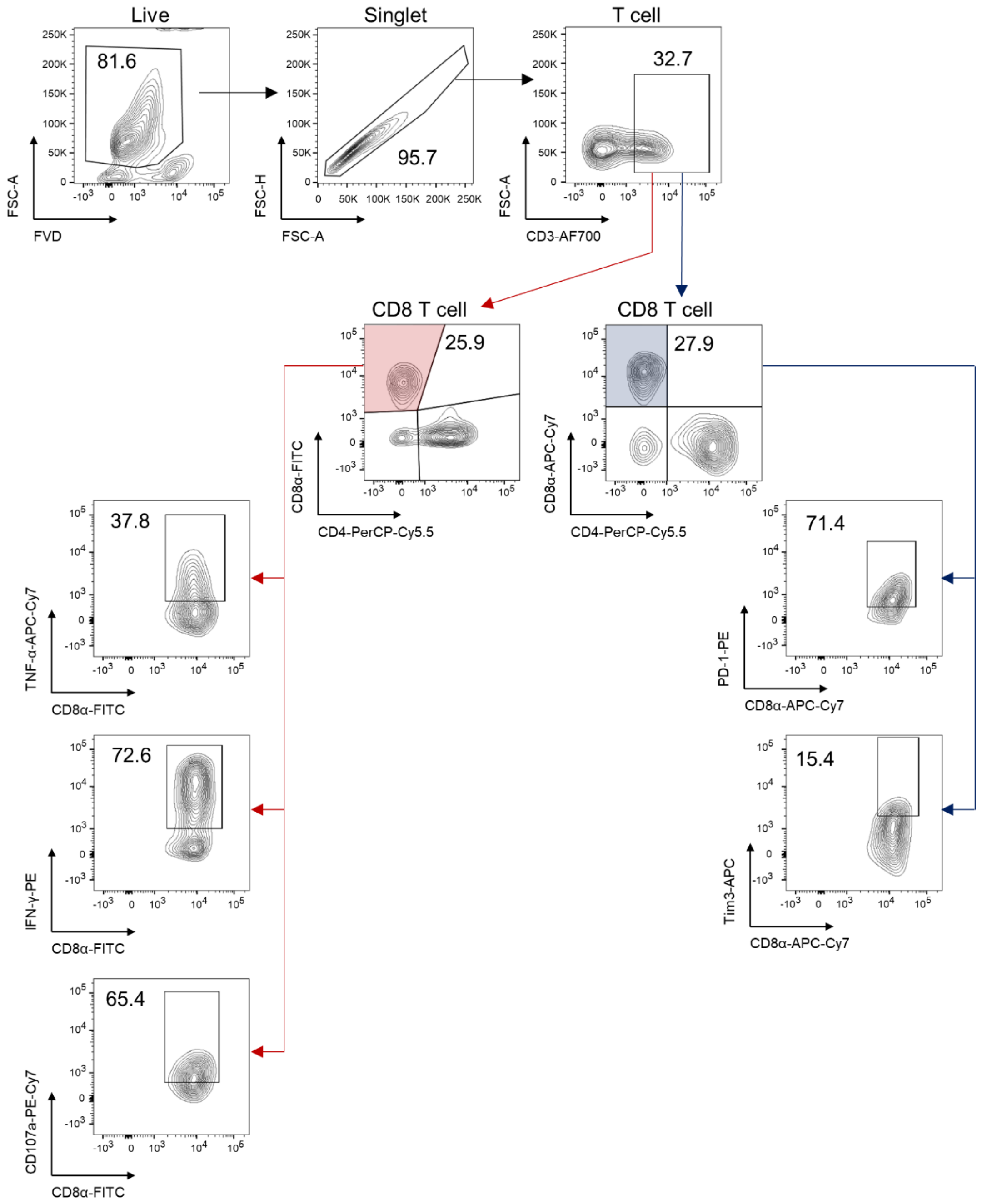


**Figure S4.** Gating strategy for the analysis of T cell effector or exhaustion molecule expression.

The gating was performed in the order depicted in the figure. The red line represents the gating flow of expression for effector molecules, and the blue line represents that of expression for exhaustion molecules.
