## Supplementary Figure S5 for "Gut microbiota dysbiosis induced by brain tumor modulates the efficacy of immunotherapy"

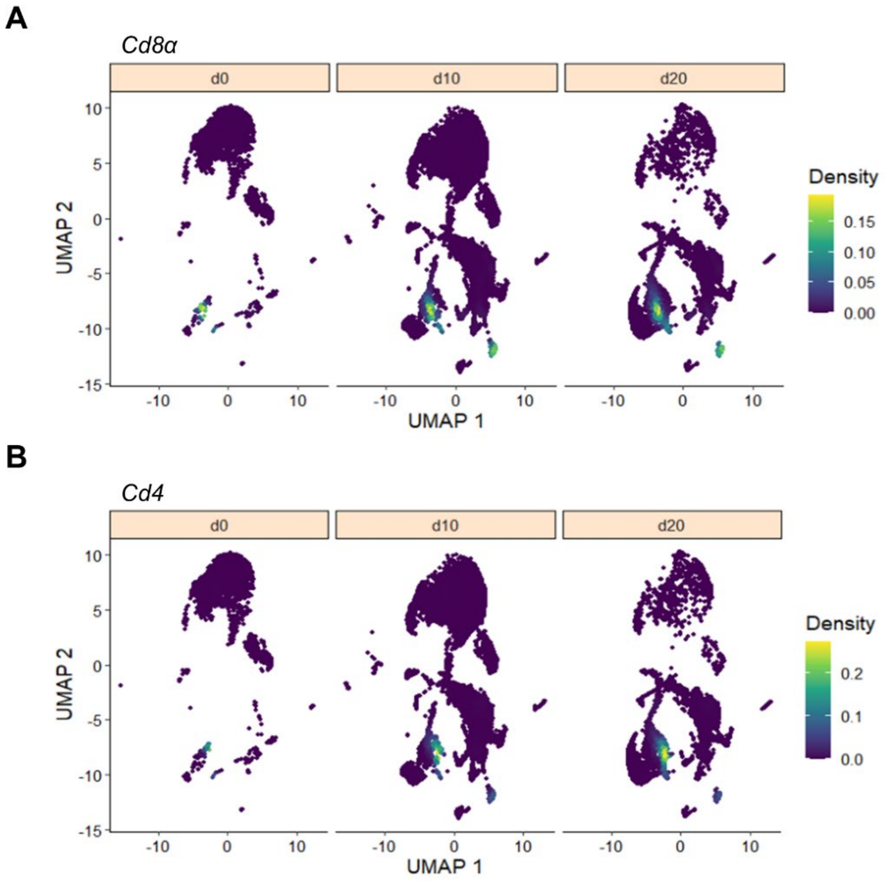


**Figure S5.** At early time points, T cells infiltrate the TME in brain tumors.

For temporal analysis, control samples and TME samples (d10 and d20) prepared D10 or D20s after tumor inoculation were integrated. The infiltration of CD8 T cells (**A**) and CD4 T cells (**B**) was assessed by the expression of *Cd8α* and *Cd4*, respectively.
