## Supplementary Figure S6 for "Gut microbiota dysbiosis induced by brain tumor modulates the efficacy of immunotherapy"

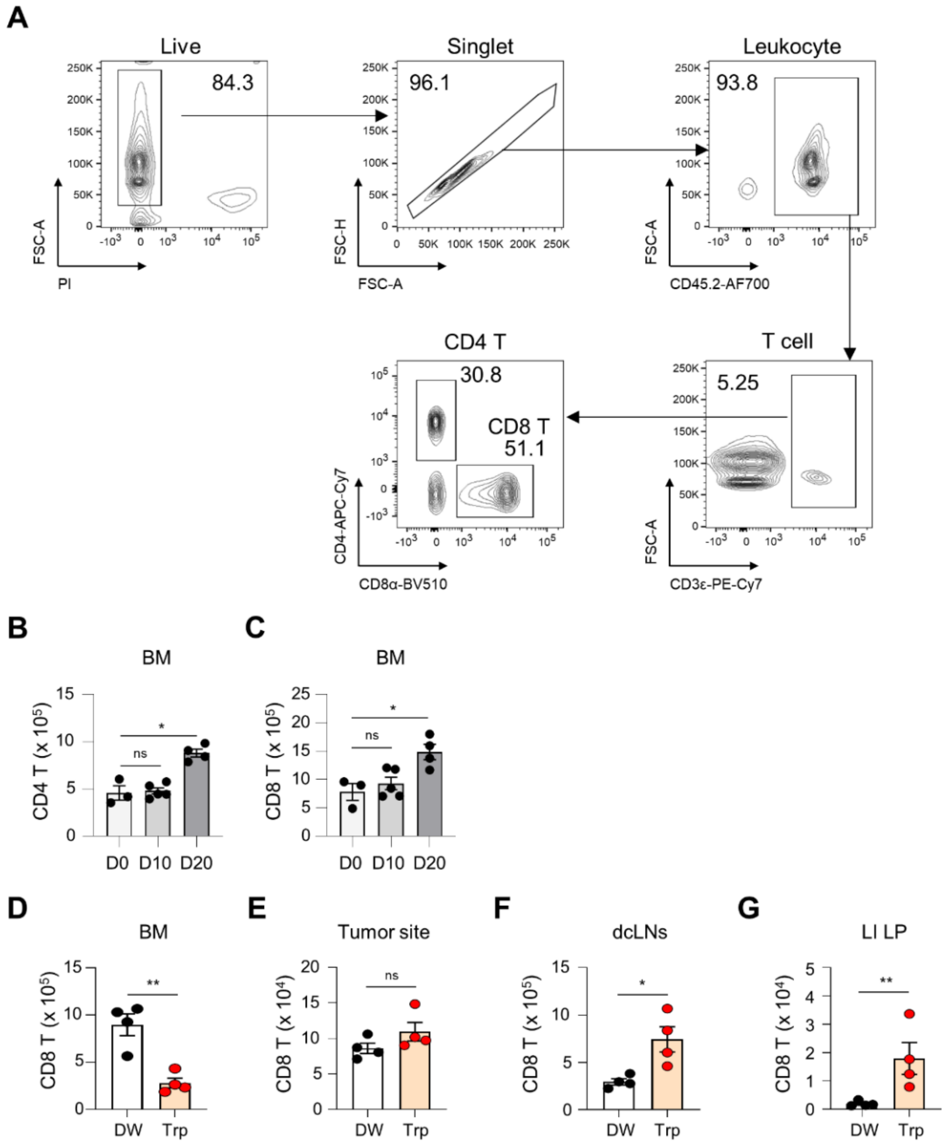


**Figure S6.** Trp supplementation enhances the circulation of CD8 T cells.

**A** Gating strategy for T-cell gating. **B**, **C** D10 (n = 5) or D20 (n = 4) after tumor implantation, bone marrow cells from tumor-bearing mice were analyzed using flow cytometry using bone marrow of healthy mice (n = 3) as the control. **D**-**G** On D20, T cells were quantified in bone marrow (**D**), tumor site (**E**), deep cervical lymph nodes (**F**), and large intestine lamina propria (**G**) by flow cytometry from control (DW, n = 4) and Trp-treated tumor-bearing mice (n = 4). We used a two-tailed, unpaired Student's *t*-test for statistical significance comparison. Data are presented as mean ± s.e.m. $*P < 0.05$, $**P < 0.01$, ns: not significant.
