## Supplementary Figure S7 for "Gut microbiota dysbiosis induced by brain tumor modulates the efficacy of immunotherapy"

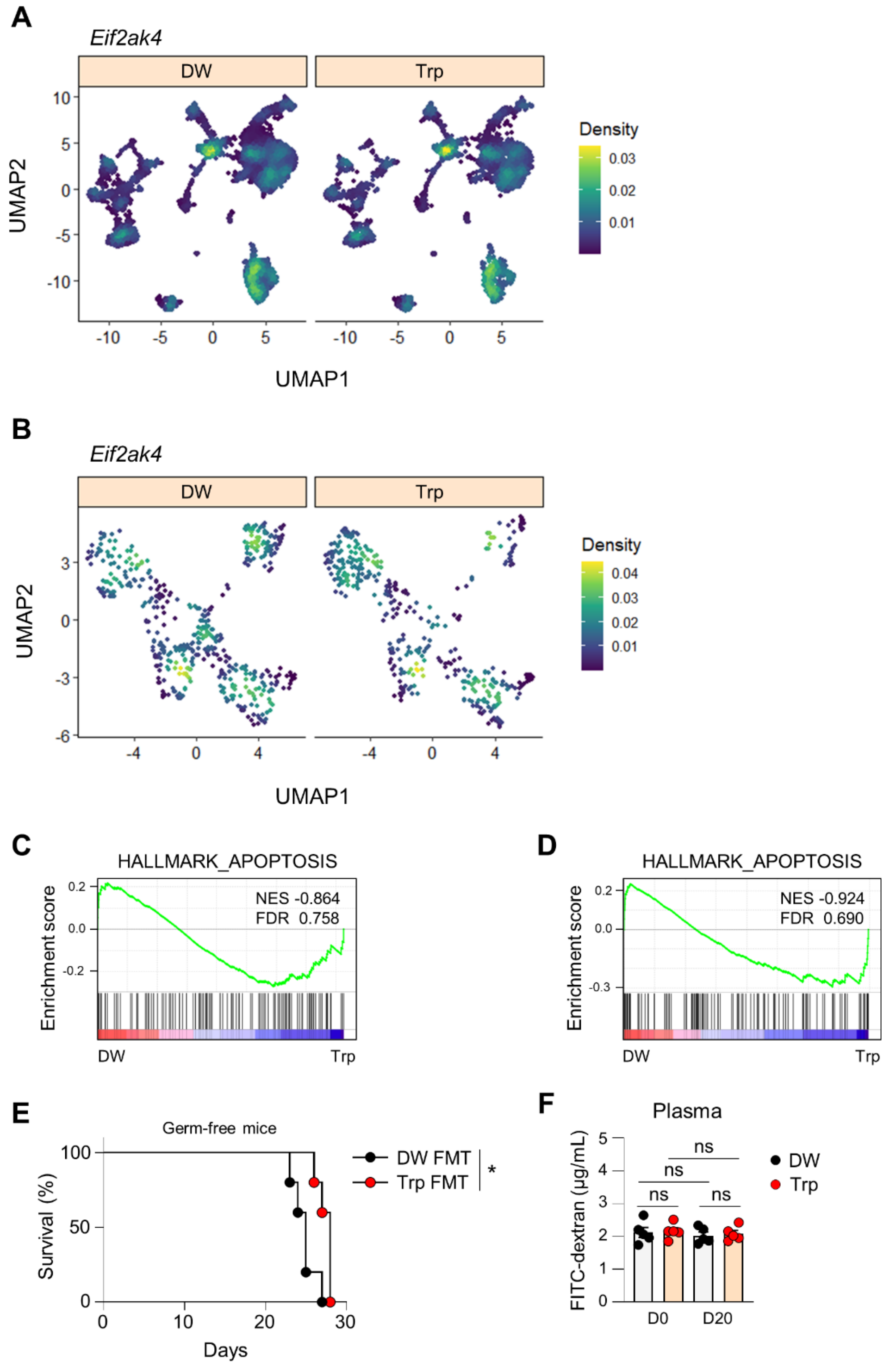


**Figure S7.** The Trp-mediated effect depends on the gut microbiota.

**A**, **B** The expression of *Eif2ak4* (GCN2) was compared in the control (DW) and Trp-treated group (Trp) for total tumor-infiltrating lymphocytes (TILs) (**A**) and CTLs (**B**). **C**, **D** GSEA analysis for HALLMARK_APOPTOSIS in T cells (**C**) and B cells (**D**). **E** Survival was determined for GF mice colonized with fecal microbiota from control (DW) or Trp-treated tumor-bearing mice (Trp) for 4 weeks before tumor cells were implanted into their brains. **F** To measure gut barrier integrity on D0 and D20 after tumor implantation, mice that had fasted for 4 hours were treated with 4 kDa dextran-FITC *per os,* and 4 hours later, fluorescence was measured in the plasma. For statistical analysis, a log-rank test was used in (**E**), and a two-tailed unpaired Student's *t*-test was used in (**F**). Data are presented as mean ± s.e.m. $^*P < 0.05$, ns: not significant.
