## Supplementary Figure S8 for "Gut microbiota dysbiosis induced by brain tumor modulates the efficacy of immunotherapy"

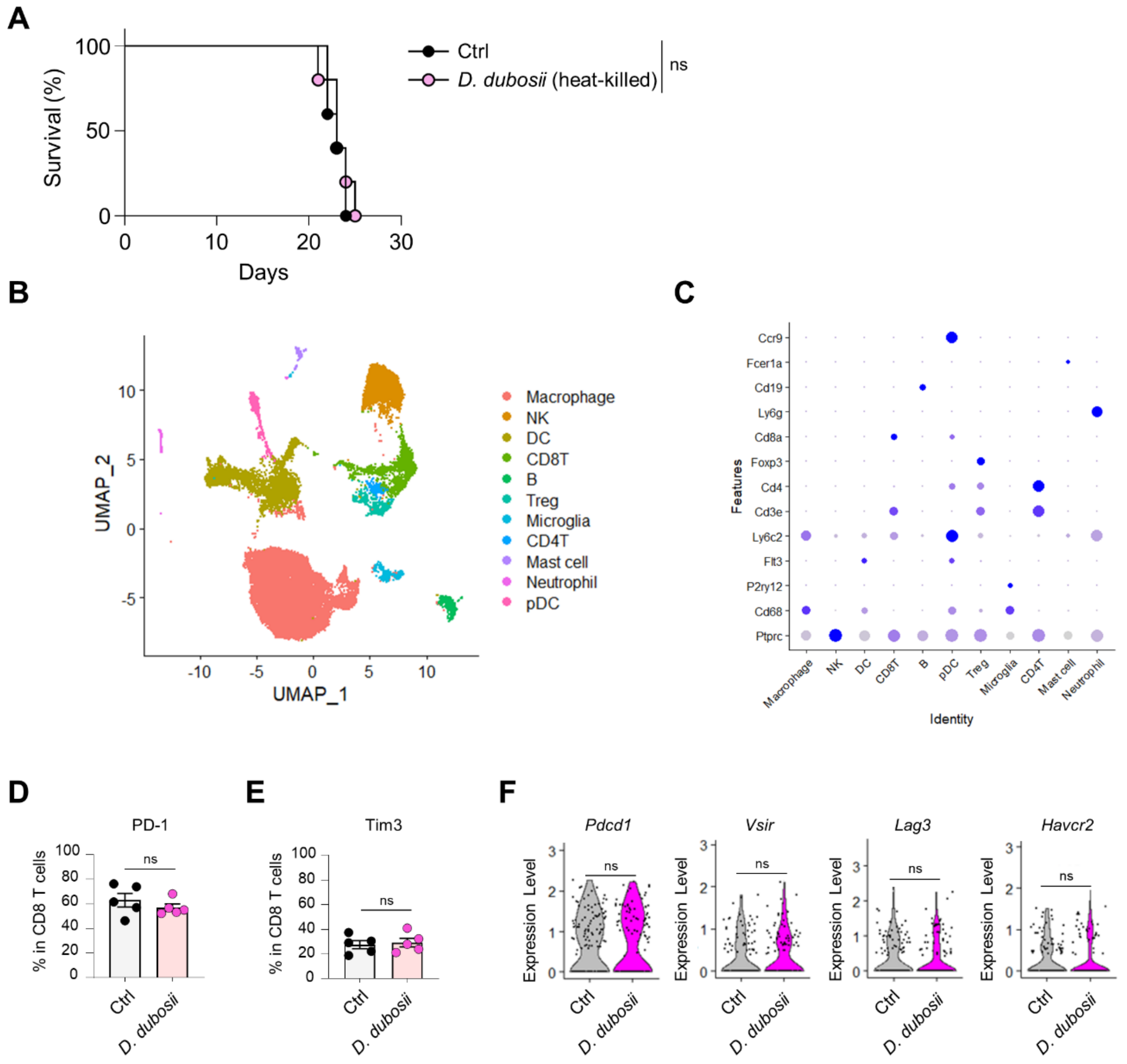


**Figure S8.** Trp-mediated changes in CTLs are observed after colonization of GF mice with *living D. dubosii*.

**A** Survival of GF mice colonized with heat-killed *D. dubosii per os* for 8 weeks, starting from 4 weeks before tumor inoculation, was measured after tumor implantation. **B**, **C** Transcriptomic analysis of tumor infiltrating leukocytes from GF tumor-bearing mice colonized or not with *D. dubosii*. **D-F** The expression of exhaustion molecules was measured using flow cytometry (**D**, **E**) and single-cell RNA sequencing (**F**). Both analysis were conducted on D20 (n =5 for each group). For statistical analysis, a log-rank test was used in (**A**), a two-tailed unpaired Student's *t*-test was used in (**D-F**). Data are presented as mean ± s.e.m. ns: not significant.
