## Supplementary Table S1 for "Gut microbiota dysbiosis induced by brain tumor modulates the efficacy of immunotherapy"

**Table S1.** Lists for flow cytometric antibody

| Name | Clone | Color | Company | Catalog number |
|---|---|---|---|---|
| CD3ε | 145-2C11 | AF700 | Biolegend | 100216 |
| | | PE-Cy7 | BD Biosciences | 552774 |
| CD4 | RM4.5 | PE-Cy7 | BD Biosciences | 552775 |
| | | PerCP-Cy5.5 | BD Biosciences | 550954 |
| | | BV650 | Biolegend | 100555 |
| CD8α | 53-6.7 | APC | BD Biosciences | 553035 |
| | | BV510 | BD Biosciences | 563068 |
| | | PerCP-Cy5.5 | BD Biosciences | 551162 |
| | | APC-Cy7 | Biolegend | 100714 |
| | | FITC | Biolegend | 100706 |
| CD45.2 | 104 | PE | BD Biosciences | 12-0454-82 |
| | | AF700 | Biolegend | 109822 |
| CD107a | ID4B | PE-Cy7 | BD Biosciences | 560647 |
| IFN-γ | XMG1.2 | PE | Biolegend | 505808 |
| TNF-α | MP6-XT22 | APC-Cy7 | Biolegend | 506343 |
| PD-1 | J43 | PE | BD Biosciences | 561788 |
| TIM-3 | RMT3-23 | APC | Biolegend | 119706 |
| 7-AAD | | | BD Biosciences | 51-68981E |
| Fixable viability Dye eFluor 450 | | | Thermo Fisher Scientific | 65-0863-14 |
| Propidium iodide | | | Invitrogen | P1304MP |
